# Use of OWL within the Gene Ontology

**DOI:** 10.1101/010090

**Authors:** Christopher J Mungall, Heiko Dietze, David Osumi-Sutherland

## Abstract

The Gene Ontology (GO) is a ubiquitous tool in biological data analysis, and is one of the most well-known ontologies, in or outside the life sciences. Commonly conceived of as a simple terminology structured as a directed acyclic graph, the GO is actually well-axiomatized in OWL and is highly dependent on the OWL tool stack. Here we outline some of the lesser known features of the GO, describe the GO development process, and our prognosis for future development in terms of the OWL representation.

## 1 Introduction

The Gene Ontology (GO) is a bioinformatics resource for describing the roles genes play in the life of an organism, covering a variety of species from humans to bacteria and viruses[1].

The way the GO is most commonly presented in publications elides much of the underlying axiomatization and formal semantics. The most common conception is a Directed Acyclic Graph (DAG) *G* =*< V*, *E >*, where each vertex in V is a particular gene “descriptor", and E is a set of labeled edges connecting two vertexes in V. The GO is rarely used in isolation - the value comes in how databases use the GO to “annotate”^1^ genes and molecular entities. A database here can be minimally conceived of as *D* =*< A*, *M >* where M is a set of molecular entities (e.g. genes or the products of genes) and A is a set of associations where each association connects an element of V with an element of M. Currently GO has some 40k vertexes, 100k edges, and the combined set of databases using GO have 27 million associations covering 4 million genes in 470 thousand different species[3].

The users of the GO apply it in a number of ways. The simplest way is to interrogate a database, for example to find out what a gene does, or to find the set of genes that do a particular thing (the latter query making use of the edges in *E*). One of the most common uses is to find a functional interpretation of a set of genes, a so-called enrichment test. For example, given a set of genes that are active in a particular type of cancer, what are the GO classes that are statistically over represented in the description of these genes? Another use is as a component of a diagnostic tool for finding causative genes in rare diseases[17].

The simple graph-theoretic view of the GO is effective and popular, but does not take into account that GO has been enriched by an ever increasing number of OWL constructs over the years.

## 2 The Axiomatic Structure of the GO

The GO consists of over 40,000 classes, but also includes an import chain that brings in an additional 10,000 classes from 8 additional ontologies. The majority of the axioms in this import chain are within the EL++ profile, allowing for the use of faster reasoners.

For release purposes, the GO is available as a limited “standard edition” which excludes imports and external ontologies and a complete edition called *go-plus*^2^. There are additional experimental extensions which are not discussed here (but we encourage reasoner developers to contact us for access to these for testing purposes).

Table 1 shows the breakdown of axiom types and expression types. As is evident, existential restrictions and intersections are frequently used, with the latter used entirely within equivalence axioms.

**Table 1.** Axiom or expression types used in the GO

| Construct | Usage count |
| --- | --- |
| Axiom | 556088 |
| EquivalentClasses | 27108 |
| IntersectionOf | 27108 |
| SubClassOf | 106063 |
| AnnotationAssertion | 370394 |
| DisjointClasses | 127 |
| SomeValuesFrom | 36125 |

The part of GO that is most typically exposed to users are the SubClassOf axioms^3^, together with annotation assertions, which is weak in terms of expressivity but delivers the query abilities required by most users.

The entire ontology reasons in seconds in Elk[9], and 10 minutes in Hermit (on a standard laptop or workstation).

### 2.1 Equivalence axioms in GO

The meat and potatoes for reasoning in GO are the equivalence axioms, most typically of a “genus-differentia” form, i.e. *X EquivalentTo G and R*_1_ *some Y*_1_ *and R_n_ some Y_n_*. These were historically referred to in GO as ‘cross products’ [12], since the set of such defined classes *X* are a subset of the cross-product of the set *G* and the sets *Y*.

The existence of these axioms allow us to use reasoners to automatically classify the GO, something that is vitally important in an ontology with such a large number of classes. In the current release version of GO, over 32,000 SubClassOf axioms were inferred by reasoning, representing a substantial efficiency gain for ontology developers.

### 2.2 Inter-ontology axioms

The subset of GO most commonly exposed comprises the so-called “intra-ontology” axioms, but GO also contains a rich set of inter-ontology axioms, leveraging external ontologies (The plant ontology, Uberon, the Cell type ontology, and the CHEBI ontology of chemical entities).

The primary use case for the inter-ontology axioms is to allow for modularized development and to automatically infer the GO hierarchy. Additionally, inter-ontology axioms have the added benefit of connecting different ontologies used for classification of different types of data.

The inter-ontology axioms are present in the go-plus edition of GO, but not the core version. To avoid importing large external ontologies in their entirety, we build “import modules” using the OWL API Syntactic Locality Module Extractor.

### 2.3 Relations in the GO

We use a number of different Object Properties in these axioms, taken from the OBO Relations Ontology^4^. We rely heavily on Transitivity, *SubPropertyOf* and *ObjectPropertyChain* expressions. We have recently started using the inverse of the partOf object property in some existential restrictions[2]; Elk ignores the InverseProperties axiom, which is generally not an issue, but there have been cases of errors we have only managed to detect using HermiT.

### 2.4 Constraints in the GO

As well as automatic classification, we also make extensive use of reasoning as part of our quality control pipeline, both for ontology validation, and for the validation of data about genes coming from external databases.

We encode the majority of constraints in GO as disjointness axioms. Domain and range constraints on object properties play less of a part. We achieve more powerful contextual domain-range type assertions using disjointness axioms. For example, the ‘part of’ relation is flexible regarding whether is it used between two processes (such as those found in the GO ‘biological process’ branch) or between material entities (for example, a GO subcellular component, such as synapse). This generality limits the utility of domain and range. However, the RO includes axioms of the form: ‘part of’ process DisjointWith ‘part of’ continuant Which prohibits category-crossing uses of ‘part of’ which would be invalid.

Disjointness axioms are also used in the traditional way, between siblings in a taxonomic classification, although these are typically under-specified.

We frequently have need to encode spatial and spatiotemporal constraints. For example, most of the cells in a complex organism such as yourself consist of a number of compartments, including the nucleus (the central HQ, where most of your genes live) and the cytosol (a kind of soup full of molecular machines doing their business). It is not enough to simply state that the cell and cytosol are disjoint classes. We also want to encode *spatial disjointness*, i.e. at no time^5^ do they share parts (made impossible by the existence of a membrane barrier between the two). We do this using General Class Inclusion axioms (GCI axioms), e.g.

~~~
(‘part of’ some cytosol) DisjointWith (‘part of’ some nucleus)
~~~

In some cases we can structurally simplify the axiom by using equivalent named classes such as ‘cytosolic part’ and ‘nuclear part’.

Another common type of constraint in the GO are so-called ‘taxon constraints’ [5]. The basic idea here is that the GO covers biology for all domains of life, from single-celled organisms to humans. However, many of the classes are applicable to specific lineages. For example, in describing the function of genes in a poriferan (sponge), it would be a mistake to use the GO class *brain development*, or any of its descendant classes, as these simple organisms lack a nervous system of any type. Whilst we would hope a human curator would not make such an error, the same cannot be said for algorithmic prediction methods that make use of the ‘ortholog conjecture’[18] to infer the function of a gene in one species based on the function of the equivalent gene in another species. Sponges have many of the same genes found in other animals that form synapses in the nervous system (the jury is out on whether this is a case of evolutionary loss or a case of co-option). Here it is useful to have a knowledge-based approach to validation of computational predictions.

The most obvious way to encode taxon constraints such as “nucleus part of ONLY Eukaryotic organisms” is using universal restrictions and complementation expressions, and for constraints such as “photosynthesis occurs in only NON-mammals” is to use ComplementOf expressions; however, this has the disadvantage of being outside EL++. We instead encode taxon constraints using shortcut relations[14] and disjointness axioms[11].

One place where we use UnionOf constructs is in the GO-specific extensions of the taxonomy ontology where we create grouping classes. For example, the grouping “Prokaryota” would not be in the taxonomy ontology as it constitutes a paraphyletic group - nevertheless it is useful to refer to these groups, so we create these as union classes (in this case, equivalent to the union of “Eubacteria” and “Archaea”). The increase in expressivity beyond EL++ is not a practical issue here as the groupings do not change frequently so we pre-reason with HermiT and assert the direct subclass inferences (here between Prokaryotes and the class for cellular organisms).

### 2.5 Annotation axioms

In addition to the logic axioms described above, GO makes heavy use of annotation assertion axioms, as the textual component of GO is important to our users. In particular, textual definitions, comments and synonyms (in addition to labels) are the annotation properties we use most commonly.

One of main factors that allowed us to move to OWL was the introduction of axiom annotations. We attempt to track provenance on a per-axiom level, so this feature is vital to us.

## 3 The GO Development environment

### 3.1 Transitioning to OWL

The GO was not born as a Description Logic ontology. In order to be able to take advantage of automated reasoning, it was necessary for us to *retrospectively* go back and assign equivalence axioms and other OWL axioms to existing classes, some of which date back to the inception of the project. This is in contrast to ontologies “born” as OWL following the Rector Normalization pattern[16] in which classes are *prospectively* axiomatized, at the time of creation.

The process of retrospective axiomatization was assisted in part by the detailed design pattern documentation maintained by the GO editors – see for example the documentation on developmental processes^6^. This made it possible to use lexical patterns to derive equivalence axioms[12]. For example, if a class *C* has a label “X differentiation” then we derived an axiom *C EquivalentTo ‘cell differentiation’ and results_in_acquisition_of_features_of some X*. X is assumed to come from the OBO cell type ontology, and if no such X exists then we add this.

However, the axiomatization process frequently revealed cryptic inconsistencies and incoherencies, both within the GO, and between the GO and other ontologies. Some of these are trivial to resolve, whereas others require a massive conceptual alignment of two domains of knowledge. One such case was the alignment of a biology-oriented view of metabolic processes with a chemistry-oriented view[7].

### 3.2 Editing tools

The GO development environment is a hybrid of different tools and technologies.

In the past, the GO developers exclusively used OBO-Edit [4] for construction and maintenance of the ontology. OBO-Edit only supports a subset of OBO-Format, which corresponds roughly to EL++, with the addition of other restrictions, such as limited ability to nest class expressions. However, the main limitation of OBO-Edit is the lack of integration with OWL Reasoners such as Elk.

Protege represents a superior environment for logic-based ontology development, but unfortunately lacks much of the functionality that makes OBO-Edit a productive and intuitive tool for the GO developers. These features include powerful search and rendering, visualization, inclusion of existential restrictions in hierarchical browsing, and annotation editing customized for our annotation property vocabulary.

To overcome this we have been moving to a hybrid editing environment, whereby developers use a mixture of OBO-Edit and Protege. The source ontology remains in OBO-Format, with the developers using OWLTools^7^ to perform the conversion to OWL and back. Developers are careful to remain within the OBO subset of OWL.

At first we employed this hybrid strategy tentatively, with the developers using Protege primarily as a debugging tool (for example, explanation of inferences leading to unsatisfiable classes). However, developers are gradually embracing Protege for other parts of the ontology development cycle, such as full-blown editing.

To facilitate this transition, we have been working with other software developers to create plugins that emulate certain aspects of the OBO-Edit experience. These include an annotation viewer and editor^8^, a plugin that manages the obsoletion of classes according to GO lifecycle policy^9^, and a partial port of the OE graph viewer^10^ (see figure 1).

**Fig. 1.**
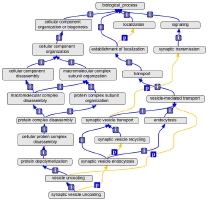
Figure 1: Protege GraphView plugin, ported from OBO-Edit

### 3.3 Web based templated term submission

Biological data curators frequently need new classes for describing the genes they are annotating. Often these classes fall into particular compositional patterns with placement in the subsumption hierarchy calculated automatically. In the past the sole method for data curators to obtain new classes was through a sourceforge issue tracking system, leading to bottlenecks.

To address this we created TermGenie[6]^11^, a web-based class submission system that allows curators to generate new classes instantaneously, provided they pass a suite of logical, lexical and structural checks. TermGenie submission can be according to either pre-specified templates, or “free-form” submissions.

Currently we specify the templates procedurally as javascript code, and we are currently exploring the use of Tawny-OWL[10] as the templating engine.

### 3.4 Smuggling OWL expressions into Databases

In order to avoid overloading the ontology with too many named classes, we have created an “annotation extension” system whereby data curators can composed their own class expressions for describing genes[8]. The expressivity of the system is deliberately limited to refining a base class using one or more existential restrictions.

This system has so far appeared to be a useful balance between expressivity and simplicity. One problem is that data curation takes place outside an OWL environment, so any logical errors (for example, violation of a domain or range constraint) are not caught until the curators submit their data to the central GO database, where we perform reasoner-based validation.

### 3.5 Ontology build pipeline

As the GO evolved from being a single standalone artifact to modular entity with a number of derived products we constructed an ontology verification and publishing pipeline.

As is common in the bioinformatics world, we specify and execute our pipeline using UNIX Makefiles, which allows the chaining together of dependent tasks that consume and produce files.

We developed a command line utility that acts as a kind OWL Swiss-army knife, with the original name of OWLTools. We developed OWLTools according to the UNIX philosophy, with a view to integration with Makefile-type pipelines. It is primarily a simple wrapper onto the OWL API, and allows the execution of tasks such as checking if an ontology is incoherent, generating ontology subsets and so on.

This pipeline is executed within a Continuous Integration framework[13].

### 3.6 Challenges of working with multiple ontologies

We aim to follow the Rector Normalization pattern, avoiding manual assertion of poly-hierarchies, instead leveraging modular hierarchies. Often these hierarchies fall in the domain of an ontology external to GO, which presents a number of challenges. This was one of the original motivations for the creation of the Open Biological Ontologies (OBO) library, to lower the barrier for interoperation, by ensuring all federated ontologies were open, orthogonal and responsive to any requirements for improvement or change. Even with these barriers lowered, challenges remain. Ontologies developed by different groups often reflect different perspectives, design patterns and hidden assumptions that can be hard to reconcile. The initial axiomatization of one ontology using another often reveals multiple unsatisfiable classes and invalid inferences. This can be time-consuming to repair. The key here is early, prospective integration, rather than after-the-fact.

There is also a deficit of tooling to support working in a multi-ontology environment. Naive construction of import chains results in highly inefficient transfer of large RDF/XML files over the web. The resulting infrastructure is fragile, with multiple points of failure. Versioning becomes of paramount importance, because simple changes in an imported ontology can wreak havoc, causing mass unsatisfiability, or loss of crucial inferences. Multiple partial solutions exist, but none are perfect. BioPortal provides URLs for individual versions of ontologies, but require an API key, which does not work well with owl imports.

The parallels with software development are obvious. As ontologies such as GO move from being monolithic to modular, we need the equivalent of dependency management and build tools such as Maven, and we welcome efforts such as the recent OntoMaven project[15].

Biological ontologies do not always modularize as cleanly as software libraries. For example, there are multiple mutual dependencies between the cell type ontology and GO (the former relies on the latter to describe what cells have evolved to do, the latter relies on the former to describe the development of these cells). This presents additional challenges.

## 4 Future developments

### 4.1 Getting OWL into the mainstream

We are highly dependent on OWL as part of the ontology development cycle in GO. Axiomatization using equivalence and disjointness axioms are crucial for automating classification and quality control. However, OWL axioms currently play less of a role once the ontology is deployed and used. There are large numbers of highly sophisticated analysis tools that incorporate the GO - most of these just treat the ontology as a simple DAG (in fact a considerable number of tools drop even this limited axiomatization, and just use the GO as a flat list of terms). We believe there is a missed opportunity here. Some of this may be in part due to the high barrier of entry for using OWL (e.g. lack of native API in languages frequently used by bioinformaticians, such as Python). This may also represent an research opportunity for algorithms that combine the types of statistical and probabilistic reasoning common in biology with powerful deductive reasoning offered by description logics.

### 4.2 Rise of the ABox

Most of the detailed axiomatization in the GO represents the low hanging fruit, such as equivalence axioms for compositional concepts. Other aspects of biology are more resistant to axiomatization - for example, multi-step pathways or complex cellular processes such as apoptosis. The tree-model property of TBoxes makes it impossible to faithfully encode multiply connected mechanistic processes. The existence of high degrees of variability and exceptions across different species presents challenges for a monotonic logical encoding.

One approach we are exploring here is to utilize the ABox to represent “prototypical” biological processes. We have developed a web-based graphical ABox editor called Noctua^12^ for exploration of this paradigm. Whilst the use of an ABox to represent knowledge lessens the power of our model for deductive inferences, it represents an opportunity for exploration of other models of inference that may be more appropriate for biological systems.

## 5 Conclusions

The OWL language and tools have benefited the GO tremendously, particularly the ontology development cycle and the validation of data about genes. At this time, GO primarily uses a subset of OWL that is supported by Elk, providing us the benefits of fast reasoning. In general, like many biological ontologies, the GO is not in need of esoteric extensions to OWL, but would benefit substantially from the creation of new tools and the hardening of existing tools, particularly related to release management.

## Footnotes

1 Note the different usage of the term *annotation* in the biological data curation world

2 http://geneontology.org/page/download-ontology

3 including existential restrictions, e.g. part_of

4 http://code.google.com/p/obo-relations

5 we elisde for now any discussion of encoding of temporal parameters in OWL

6 http://www.geneontology.org/page/development

7 http://code.google.com/p/owltools

8 https://github.com/hdietze/protege-obo-plugins

9 https://github.com/balhoff/obo-actions/downloads

10 https://code.google.com/p/obographview/

11 http://termgenie.org

12 https://github.com/kltm/go-mme

